# Glycine receptors are not directly modulated by glutamate, AP5 or NMDA

**DOI:** 10.1101/2020.03.08.982900

**Authors:** Karin Aubrey, Diba Sheipouri, Robert Vandenberg, Yo Otsu

## Abstract

3.

Reproducibility of research data is a significant problem with more than 60% of biological and medical researchers reporting they have failed to reproduce published data. General acceptance of incorrect results can mean that future data is incorrectly interpreted and progress significantly interrupted. Thus, replication studies play an essential role in corroborating research findings and validating future research objectives. Here, we attempted to replicate data demonstrating the neurotransmitter glutamate, as well as NMDA and AP5, acts as positive allosteric modulators of the inhibitory glycine receptor. Notably, it was shown that the amplitude of miniature glycinergic currents recorded in spinal cord slices were reversibly enhanced when extracellular glutamate concentrations were increased by the glutamate transporter antagonist TBOA. This finding indicates that endogenous fluctuations in extracellular [glutamate] permits cross-talk between excitatory and inhibitory synapses and likely plays a role in setting the spinal inhibitory glycinergic tone and modulating baseline neurotransmission. We re-evaluated the data in primary cultured spinal cord neurons, spinal cord slice and *Xenopus laevis* oocytes expressing recombinant glycine receptors. Despite extensive efforts, we were unable to reproduce the finding that glutamate, AP5 or NMDA positively modulate glycine receptor currents. We paid careful attention to key aspects of the original study design, ensured rapid drug exposure by using fast-flow application and took into account receptor saturation and protocol deviations such as animal species. This study refutes the finding that glycine receptors are directly modulated by glutamate spill-over and suggests that glycinergic tone is independent of changes in excitatory activity.

**Significance Statement:** Glutamate spill-over onto inhibitory synapses has been reported to positively modulate glycine receptors and alter the inhibitory tone of the spinal cord. This finding has important implications for baseline spinal transmission and could play a role when chronic pain develops. However, we failed to replicate these results and did not observe any modulation of native or recombinant glycine receptor-mediated currents by AP5, NMDA or glutamate. This indicates that inhibitory glycine receptors operate independently of fluctuations in extracellular [glutamate].

**Visual Abstract:** N/A

## 6. Introduction

In the adult spinal cord and brainstem neurons are inhibited by synaptically released glycine and/or GABA acting at functionally distinct postsynaptic glycine- or GABA_A_-receptors (Anderson et al., 2009; Foster et al., 2015; Harvey et al., 2004; Heinke et al., 2004; Kotak et al., 1998; Mitchell et al., 2007; Yevenes and Zeilhofer, 2011; Zeilhofer et al., 2012), but see (Moore and Trussell, 2017) and excited by glutamate acting at post-synaptic ionotropic AMPA- kainite- or NMDA-receptors. The balance between inhibition and excitation is essential for effective neuronal signalling and disruption of this balance is thought to contribute to neuronal diseases such as autism and chronic pain (Colloca et al., 2017; Marin, 2012).

Although each synapse generally uses a distinct fast neurotransmitter system, they do overlap in many neurons and under some circumstances can modulate each other’s activity (Ahmadi et al., 2003; Chaudhry et al., 1998; Chery and De Koninck, 1999; Lu et al., 2008; Sagne et al., 1997; Sun et al., 2014). For example, glycine is an agonist at GlyR and a co-agonist with glutamate at N-methyl-D-aspartate receptors (NMDAR) (Johnson and Ascher, 1987) and thus has both inhibitory and excitatory actions. In the spinal cord and hind-brain, synaptically released glycine can spill over from inhibitory synapses onto nearby excitatory synapses and influence NMDA-receptor mediated signalling (Ahmadi et al., 2003), this form of cross-talk is thought to play an important role in balancing inhibitory and excitatory signalling (Kullmann, 2000).

Glycine and GABA signalling also overlap pre- and post-synaptically at some inhibitory synapses. Presynaptically, GABA and glycine share a vesicular transporter (Chaudhry et al., 1998; Sagne et al., 1997) and can be co-packaged and co-released from the same synaptic vesicles (Aubrey et al., 2007; Jonas et al., 1998). GABA/glycine co-release occur at ~ 30% of inhibitory synapses in the adult hind-brain and, although postsynaptic GABA_A_R and GlyR are usually differentially expressed at the corresponding postsynaptic density, co-release is thought to influence inhibitory transmission by engaging extra-synaptic and presynaptic receptors (Chery and De Koninck, 1999; Chery and De Koninck, 2000; Keller et al., 2001; Lim et al., 2000; Turecek and Trussell, 2001). In addition, GABA can bind to postsynaptic glycine receptors and accelerate their decay kinetics (Lu et al., 2008).

In a paper published in the journal “Nature Neuroscience” in 2010, Liu and colleagues demonstrated that glutamate can spill-over onto nearby inhibitory synapses and allosterically facilitate GlyR mediated currents (max effect ~ 200%) (Liu et al., 2010). Initially, it was found that miniature glycinergic inhibitory post synaptic currents (mIPSCs) recorded from rat spinal cord cultures were enhanced when the competitive NMDAR antagonist D-(-)-2-Amino-5-phosphonopentanoic acid (AP5) was added to the extracellular bath solution. Next, the authors showed that this effect was mimicked by glutamate and a range of other glutamate receptor agonists including NMDA, kainic acid and quisqualate and the glutamate receptor antagonist kynurenic acid, but not by the pore blocking NMDAR antagonist MK-801 or competitive AMPA receptor antagonists NBXQ and CNQX. The effect was verified in HEK cells expressing recombinant GlyRs, suggesting a direct interaction of glutamate and its analogues with GlyRs and the results were strengthened by single channel recordings of native GlyR currents demonstrating that the presence of D-AP5 increased open dwell times without altering conductance levels, and that the GlyR enhancing effects were only observed if the drug was applied extracellularly. Finally, the paper showed that the amplitude of glycinergic mIPSCs recorded in rat spinal cord slices were increased by 42%, without no change in frequency, when the extracellular glutamate concentrations were elevated by the presences of the glutamate transporter antagonist TBOA. These findings are important because they suggest that the size and duration of native GlyR currents will be determined not only by the relative concentrations of glycine and GABA released from presynaptic vesicles coupled to the density of postsynaptic receptor isoforms, but also by the extent of glutamate spill-over from nearby synapses. In addition, all *in vitro* measures of GlyR currents previously reported in the presence of AP5 or kynurenic acid should be re-evaluated.

The hypothesis that GlyR and NMDAR mediated transmission is reciprocally modulated by neurotransmitter spill-over and its role in tuning pain circuits has never been investigated. We first sought to replicate the finding that AP5, NMDA and glutamate act as positive allosteric modulator (PAM) of GlyRs (Liu et al., 2010) and recorded GlyR-mediated currents from whole-cell patch clamped mouse primary cultured spinal cord neurons, dorsal horn neurons in rat spinal cord slices and from oocytes expressing recombinant GlyR. Surprisingly, we were unable to find any evidence that AP5, NMDA or glutamate facilitate glycine currents, instead we observed no change or a small reduction in current amplitude. Our findings are discussed in the context of variations in the experimental protocol and possible confounding factors.

## 7. Materials and methods

### Ethical approval

All procedures involving animals followed the guidelines of the ‘Australian Code of Practise for the Care and Use of Animals for Scientific Purposes’ and with the approval of the Royal North Shore Hospital Animal Ethics Committee (rodent work) or the University of Sydney Animal Ethics Committee (Xenopus)

Rats were housed in groups of 2-3 and mice in groups of up to 5 in individually ventilated cages under a 12:12 h light/dark cycle, with environmental enrichment and free access to water and standard rat chow.

### Recordings from neurons

#### Embryonic mouse spinal cord neurons

Primary cultures of spinal cord neurons were prepared as described by Hanus et al. (2004) (see also (Rousseau et al., 2008)) from embryonic day 13-14 C57BL/6 wild-type or GlyT2::Cre × mTmG mouse pups. Embryos were obtained by caesarean section from pregnant mice (n = 7) deeply anesthetized with isoflurane (3 %; assessed by rate of breathing, lack of righting reflexes and lack of withdrawal reflex in response to hindpaw squeeze) and killed by cervical dislocation. Spinal cords were dissected under sterile conditions into PBS with 33 mM glucose at pH 7.4 and then incubated in trypsin/EDTA solution (0.05% v/v) for 10 min at 37°C. Cells were dissociated mechanically in a modified L15 Leibowitz’s medium (Invitrogen) and plated at a density of 1.4 −2.0 × 10^5^ cells/cm^2^ on sterilized glass coverslips coated with 60 μg/ml poly-DL-ornithine and with medium containing 5% inactivated fetal calf serum (Sigma). Cells were maintained at 37°C in 5% CO2 in serum-free Neurobasal plus medium containing supplement B27 plus (Invitrogen) for up to 3 weeks. Medium was changed every 3-4 days.

*Spinal cord slices* were obtained from 2 male adult Sprague Dawley rats (8 - 10 weeks old) who had vector-mediated channel rhodopsin expressed in rostroventral medial medulla neurons. Rats were obtained from Animal Resources Centre (Canning Vale, Australia). Rats were deeply anaesthetized with isoflurane (3 %, assessed by rate of breathing, lack of righting reflexes and lack of response to paw squeeze), transcardially perfused with ice-cold NMDG solution; (in mM, 93 NMDG; 30 NaHCO_3_; 25 glucose; 2 thiourea; 3 Na-pyruvate; 2.5 KCl; 1.2 NaH2PO_4_-H2O; 20 HEPES; 10 MgSO_4_-7H2O; 0.5 CaCl_2_;5 Na-Ascorbate; ~300 mOsm, equilibrated with 95 % O_2_-5% CO_2_) and parasagittal spinal cord slices (280 μm) were prepared with a vibratome (VT1200S, Leica Microsystems AG, Wetzlar, Germany) in the same solution. After maintaining the slices for 10 minutes at 34°C in a submerged chamber, they were kept in artificial cerebrospinal fluid (ACSF) equilibrated with 95% O_2_ and 5% CO_2_ until recording. The slices were then individually transferred to a chamber and superfused continuously (2.5 ml min^−1^) with ACSF (32°C, composition (in mM):126 NaCl, 2.5 KCl, 1.4 NaH_2_PO_4_, 1.2 MgCl_2_, 2.4 CaCl_2_, 11 glucose and 25 NaHCO_3_).

#### Electrophysiology

Whole-cell voltage-clamp recordings of cultured spinal cord neurons [12–22 d in vitro (DIV)] were performed at ~32°C using a Multiclamp 700B controlled and Axograph acquisition software. Currents were filtered at 4 kHz and sampled at 20 kHz using a National Instruments USB-6251 digitizer. IPSCs were filtered (20 kHz low-pass filter) and sampled (4 kHz) for on-line and later off-line analysis (Axograph 1.7.6). A sample size of 4-6 cells per group was anticipated prior to the experiments based on previous experience and the results of Liu et al (2010). Cells where the series resistance of the recorded neuron were less than 20 MΩ and neurons where it changed by more than 25% were excluded from the analysis. Patch pipettes were pulled from borosilicate glass capillaries (Hilgenberg) and had resistances of 4-6 MΩ. Miniature and evoked currents were recorded at a holding potential (VH) of −60 or −65 mV using pipettes filled with internal solution containing the following (in mM):140 CsCl, 1 CaCl2,, 1 MgCl2, 10 EGTA, 1 BAPTA, 4 Mg-ATP, 5 QX314 [N-(2,6-dimethylphenylcarbamoylmethyl)triethylammonium-Cl], and 10 HEPES, (pH 7.4 with CsOH, 300 ± 5 mOsm. In Figures 4 and 5 the internal solution used was modified to be identical to that used in (Liu et al., 2010) and contained (in mM) 140 CsCl, 10 BAPTA, 4 Mg-ATP and 5 QX-314 and 10 HEPES, (pH 7.20, osmolality, 295 ± 5 mOsm). For paired recordings, the internal solution for the presynaptic current clamped neuron contained (in mM):155 K-gluconate, 4 KCl, 5 Mg-ATP, 0.1 EGTA, and 10 HEPES, adjusted to pH 7.4 with KOH. Neurons were continuously bathed with an external solution containing the following (mM):140 NaCl, 5.4 KCl, 10 HEPES, 1 MgCl_2_, 1.3 CaCl_2_ and 20 glucose (pH 7.4, 305-315 mOsm). AMPA and NMDA receptors were blocked with 10 μM CBQX and when indicated 15 μM MK-801. GABA_A_receptors (GABA_A_Rs) were selectively blocked with 10 μM Bicuculline. When indicated, GlyRs were blocked with 0.5-1 μM strychnine. Miniature events were recorded in the presence of 0.5 μM tetrodotoxin (TTX). A concentration of AP5 equivalent to 100 μM D-AP5 was used in all experiments (eg. 100 μM D-AP5 or 200 μM DL-AP5), NMDA was used at 50 μM and glutamate at 100 μM. Drugs were purchased from Sigma or Tocris.

### 2-electrode voltage clamp recording of heterologously expressed human glycine receptors in Xenopus laevis oocytes

Human GlyRα1 and GlyRβ cDNA were subcloned into pGEMHE. The amplified cDNA/pGEMHE product was then transformed in E. coli cells, and subsequently purified using the PureLink Quick Plasmid Miniprep Kit (Invitrogen by Life Technologies, Löhne, Germany), and sequenced by the Australian Genome Research Facility (Sydney, Australia). The purified plasmid DNA was linearized via the restriction enzyme NheI (New England Biolabs (Genesearch), Arundel, Australia). Complementary RNAs were synthesised using the mMESAGE mMACHINE T7 kit (Ambion, Texas, USA).

*Xenopus laevis* frogs (n = 2) were anesthetised with 0.17% (w/v) 3-aminobenzoic acid ethyl ester and surgical anaesthesia was assessed by a regular and relaxed respiratory rate, no withdrawal reflex when the hind feet are pinched, no muscle tone when hind limb is extended and no response to external stimuli. Then, an ovarian lobe was removed via an incision in the abdomen and stage V oocytes were isolated from the lobe via digestion with 2 mg mL-1 collagenase A (Boehringer, Mannheim, Germany) at 26°C for 1 hour. 2 ng of cRNA encoding GlyRα1 was injected into each oocyte cytoplasm when the receptors were studied individually. Where GlyRα1β were expressed, a 1:5 ratio of GlyRα1 (2 ng) and GlyRβ (10 ng) cRNA was injected into single cells, as this ratio was found to be sufficient for formation of GlyR heteromers as judged by reduced sensitivity to pictrotoxin compared to GlyRα1 homomers. The oocytes were then stored in frog Ringer’s solution (96 mM NaCl, 2 mM KCl, 1 mM MgCl2, 1.8 mM CaCl2, 5 mM HEPES, pH 7.5) which was supplemented with 2.5 mM sodium pyruvate, 0.5 mM theophylline, 50 μg/mL gentamicin and 100 μM mL-1 tetracycline. The oocytes were stored at 18°C for 2-5 days, until receptor expression was adequate for measurement using the two-electrode voltage clamp technique. Oocytes were voltage clamped at −60 mV, and whole-cell currents generated by the substrate were recorded with a Geneclamp 500 amplifier (Axon Instruments, Foster City, California, USA) and a Powerlab 2/20 chart recorder (ADInstruments, Sydney, Australia).

### Data analysis

A sample size of 4-6 cells per group was anticipated prior to the experiments based on previous experience and the results of Liu et al (2010). *mIPSCs* were detected using a template protocol (SD = −3.5) in Axograph. mIPSC currents of less than 10 pA amplitude and with rise time and decay time values outside 0.1 - 1ms and 1 - 10 ms respectively were discarded. In cultured neurons, the mIPSC peak current and frequency was determined before and after exposure to drug by averaging miniature currents recorded over 1 minute during the 1-2 minutes preceding and proceeding bath application of the drug. In 2 cases when mIPSC frequency was low (<1Hz), mIPSCs average was calculated over 2 minutes.

In rat spinal cord slices where the frequency of mIPSCs was low and solution changes took longer to have an effect, the mIPSC peak current and frequency was determined before and after exposure to drug by averaging miniature currents recorded over 2 minute preceding application of the drug and after 4 minutes of drug wash in.

In experiments using the GlyR PAM zinc, zinc was extracellularly applied at a concentration of 2 μM free zinc (200μM Zinc chloride in buffer containing 10 mM tricine (Paoletti et al., 1997)).

The current and frequency of mIPSCs were compared with a two-tailed paired t-test and the mean and 95% confidence interval of the ratio of drug-treated/control values was reported.

*Evoked IPSC* peak current and kinetic was determined before and after exposure to AP5 by averaging 5 consecutive eIPSCs currents (triggered every 30 s) preceding and proceeding bath application of the drug. eIPSCs were measured after > 1 min wash in of drugs. In half the experiments presented the NMDA receptor antagonist MK-801 (15 μM) was included in the bath solution but as the result was not altered this data was pooled. The current and kinetics of eIPSCs were compared with a two-tailed paired t-test and the mean and 95% confidence interval of the ratio of drug-treated/control values was reported.

*Exogenous glycine currents* were stimulated by extracellular 10-50 μM glycine ± drug (1-3 seconds with a fast-step system) triggered every 1 minute, to allow glycine to washout from the bath and to minimise the effects of receptor desensitization. Current amplitude were measured before and after exposure to drug (AP5, NMDA) by averaging 3 consecutive currents. The current amplitude was compared with a two-tailed paired t-test and the mean and 95% confidence interval of the ratio of drug-treated/control values was reported. Recordings where NMDA was applied were carried out in modified extracellular solution that was Ca2+ free and contained EDTA (10mM) and 15 μM MK-801 to prevent NMDAR mediated effects

*Oocytes* expressing either α1 or α1β GlyRs were placed in an oval-shaped bath with a volume of 0.5 mL, with laminar flow of frog Ringer’s solution around the oocyte at a rate of 10 mL min-1 under gravity feed. Glycine and/or glutamate was dissolved in frog Ringer’s solution and bath applied and whole-cell currents were measured. Consecutive currents were stimulated 3 minutes apart to allow glycine to washout of the bath and to minimise the effects of receptor desensitization. The peak currents measured before and after exposure to glutamate were measured and the mean and 95% confidence interval of the ratio of glutamate-treated/control values was reported.

## 8. Results

### Synaptic glycine currents were not enhanced by AP5

We recorded GlyR miniature IPSCs from cultured mouse spinal cord neurons in the presence and absence of the competitive NMDAR antagonist AP5. The presence of AP5 did not substantially alter glycine mIPSC peak amplitude or frequency (Figure 1A,C) or kinetic (Fig. 1B, n = 6 experiments had similar kinetic results). The mean amplitude in the presence of AP5 was similar to control (AP5/control ratio = 1.029; 95% confidence interval:0.917 to 1.14; P = 0.611 in two-tailed paired t-test). In contrast, free zinc (2 μM), a well-established PAM of glycine receptors (Lynch et al., 1998; Miller et al., 2005), was able to substantially increase glycine mIPSC amplitude without effecting frequency (Figure 1D,E). The mean amplitude in the presence of zinc was 1.342 times larger than control currents (95% confidence interval:1.006 to 1.679; P = 0.034 in two-tailed paired t-test), while the mean frequency remained stable (1.102 of control; confidence interval:0.8423 to 1.362; P = 0.2628 in two-tailed paired t-test).

**Figure 1:**
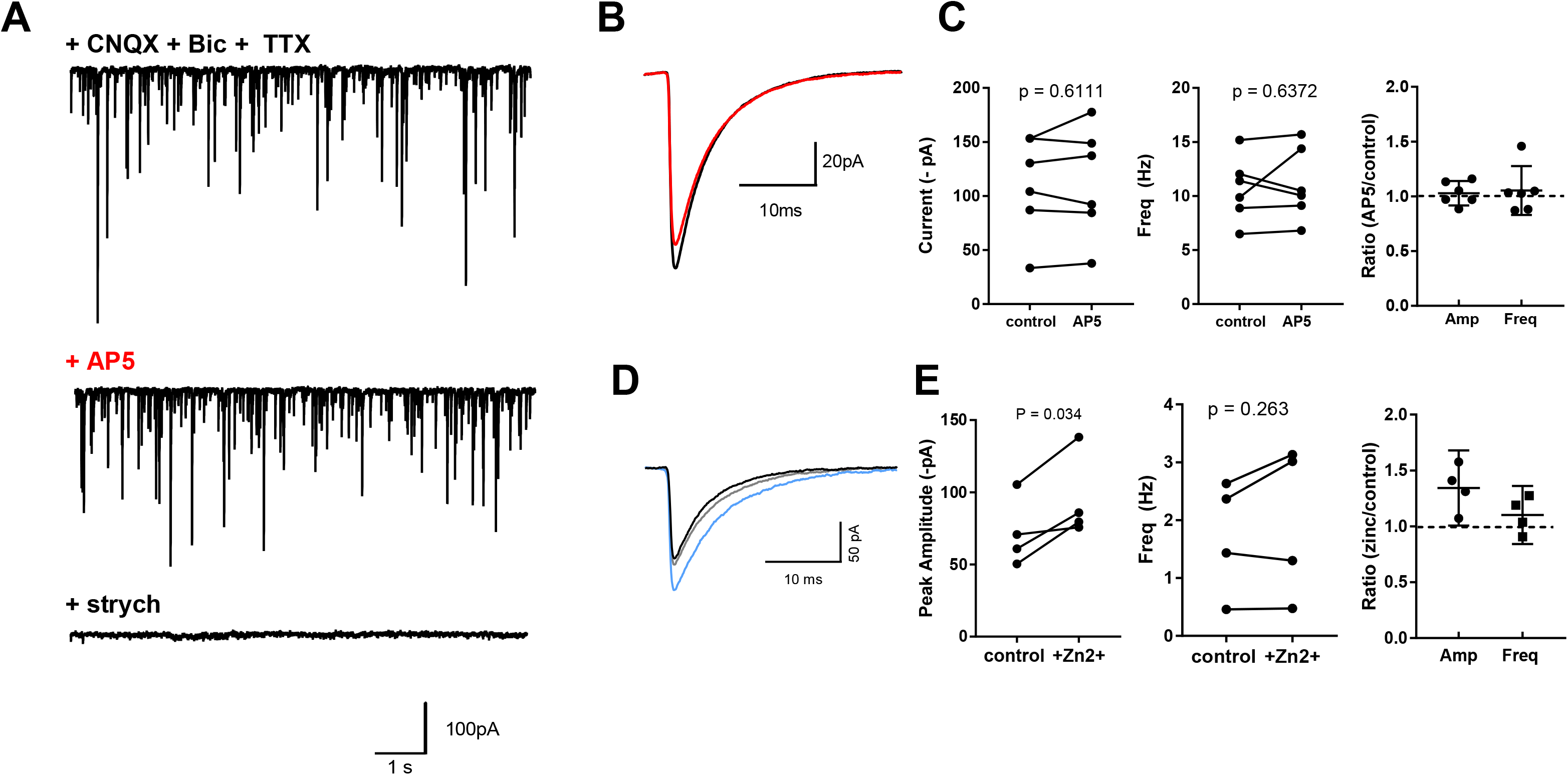
Glycinergic mIPSC recorded in cultured spinal cord neurons were not altered by AP5. A. Example traces of mIPSCs recorded in control conditions and after the addition of AP5 and strychnine to the bath solution. B. The average peak amplitude of individual experiments were compared before (black line) and after (red line) exposure to 100 μM D-AP5. C. Raw data points from individual experiments are connected by a line and proportional change (Ratio AP5/control) of mIPSC amplitude and frequency from individual experiments is shown. D. Extracellular zinc (blue line) enhanced the average peak amplitude of mIPSCs. E. The average peak glycine mIPSCs before (control) and after (Zn^2+^) exposure to 2μM free zinc. Data points from individual experiments are connected by a line and proportional change (Ratio zinc/control) of mIPSC amplitude and frequency from individual experiments are shown. The error bars shows the mean and its 95% confidence interval from n = 4-6 cells.

Similarly, when we recorded evoked glycinergic IPSCs (eIPSC) from pairs of connected neurons, the presence of AP5 did not substantially alter glycine eIPSC peak amplitude or kinetics (Figure 2). The mean amplitude in the presence of AP5 was similar to control (AP5/control ratio = 0.906; 95% confidence interval:0.6729 to 1.139; P = 0.7255 in two-tailed paired t-test).

**Figure 2:**
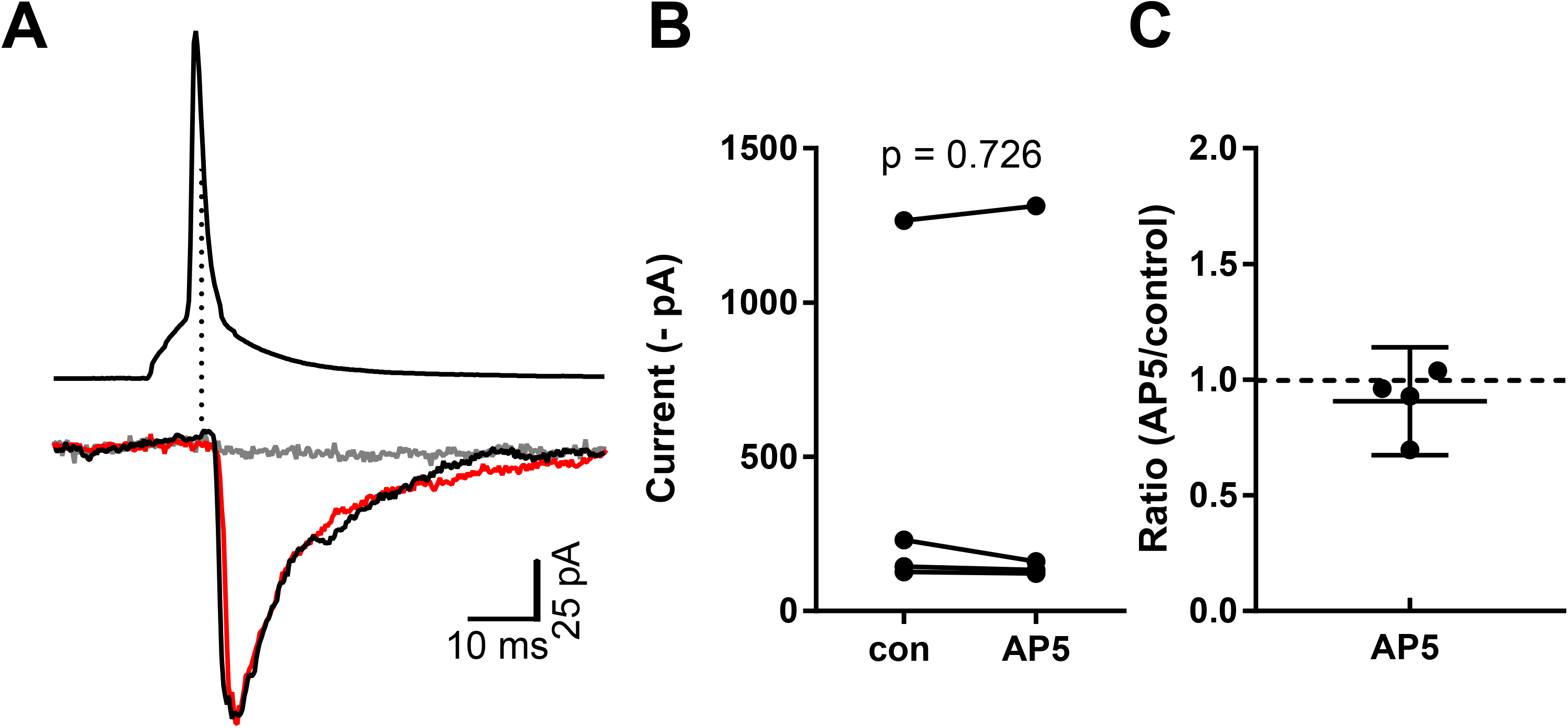
A. The average peak amplitude of individual experiments were compared before (con) and after (AP5) exposure to 100 μM D-AP5. B. Data points from individual experiments are connected by a line and proportional change (Ratio AP5/control) of eIPSC amplitude and frequency from individual experiments are shown. The error bars shows the mean value and its 95% confidence interval from n = 4 cells.

### Extrasynaptic receptors are not enhanced by AP5 or NMDA

In contrast to synaptic m/eIPSCs, short application of extracellular glycine activates extra-synaptic as well as synaptic receptors. To test the hypothesis that AP5 may modulate extrasynaptic forms of the GlyR, which are likely to be constructed from a more diverse range of GlyR subunits and interact with distinct partner proteins (Triller and Choquet, 2005), we recorded GlyR currents evoked by the extracellular application of glycine in the presence and absence of AP5 or the selective NMDAR agonist, NMDA (Figure 3) that was also reported to effectively potentiate glycine receptor currents (Liu et al., 2010). These recordings were carried out in modified extracellular solution (Ca^2+^ free with EDTA and MK-801) to prevent NMDAR mediated effects.

**Figure 3:**
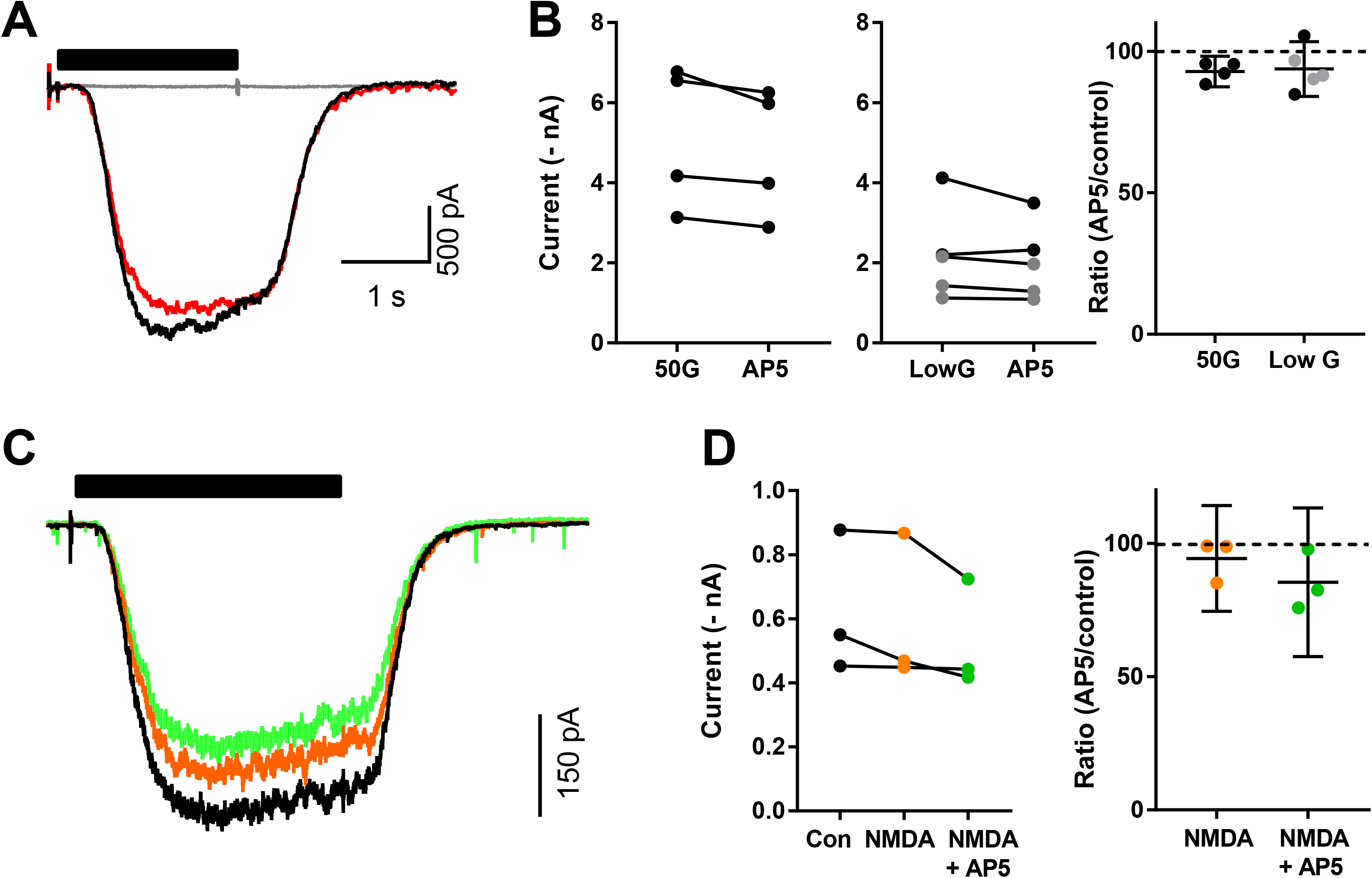
A. Exogenous glycine currents stimulated with 10 μM glycine before (black line) and after exposure to 100 μM AP5 (A, red; and with strychnine, grey), or with 50 μM NMDA and then NMDA + AP5 (C, orange and green lines respectively). Individual glycine current amplitudes stimulated with 50 μM glycine (B, left) or with low concentration glycine (B, middle; 10 μM, light circles or 30 μM, dark circles) before and after exposure to AP5. Data points from individual experiments are connected by a line and proportional changes (Ratio drug/control) of glycine currents from individual experiment are shown. The error bars shows the mean and its 95% confidence interval from n = 4-5 cells.

Glycine receptor mediated currents were stimulated by extracellular application of glycine (10-50 μM) in the presence and absence of AP5, NMDA or co-application of NMDA + APV. The mean amplitude of the whole-cell current measured was not enhanced by the presence of AP5 alone (Figure 3A,B) nor by the presence of NMDA-alone or NMDA+AP5 (Figure 3CD; current in NMDA, 0.943 of control, confidence interval:0.745 to 1.141; current in NMDA+AP5, 0.854 of control; confidence interval:0.575 to 1.133; adjusted P = 0.1589, repeat measure one-way ANOVA). In addition, while the whole-cell current amplitude was reduced by approximately half when the glycine concentration was lowered from 50 μM to low glycine (10-30 μM, Figure 3B), no AP5 effect was uncovered (50G + AP5, 0.929 of control; confidence interval:0.875 to 0.983; P = 0.0708 n two-tailed paired t-test; LowG + AP5, 0.934 of control; confidence interval:0.834 to 1.034; P = 0.2354 in two-tailed paired t-test)

### Intracellular solution and species difference do not account for the lack of effect

Intracellular Ca^2+^ transients are another positive modulator of glycine IPSCs (Fucile et al., 2000; Lévi et al., 2008; Xu et al., 2000). Our intracellular solution had slightly different calcium buffering capacity then the intracellular solution used in the original study (1 mM Ca^2+^+ 1 mM BAPTA + 10 mM EGTA compared to 10 mM BAPTA). To rule out the possibility that the intracellular solution used in (Liu et al., 2010) allowed fluctuations in intracellular calcium that may account for the GlyR potentiation observed, we replicated intracellular and extracellular solutions of (Liu et al., 2010) and recorded mIPSCs in the presence and absence of AP5. The mIPSC amplitude and frequency (Figure 4A,B) was not changed by AP5 when the intracellular solution was changed and included high BAPTA. The mean amplitude in the presence of AP5 was 1.059 times that of control currents (95% confidence interval:0.776 to 1.446; P = 0.481 in two-tailed paired t-test), and the mean frequency was 1.77 times that of control; confidence interval: −0.531 to 4.071; P = 0.2628 in two-tailed paired t-test).

**Figure 4:**
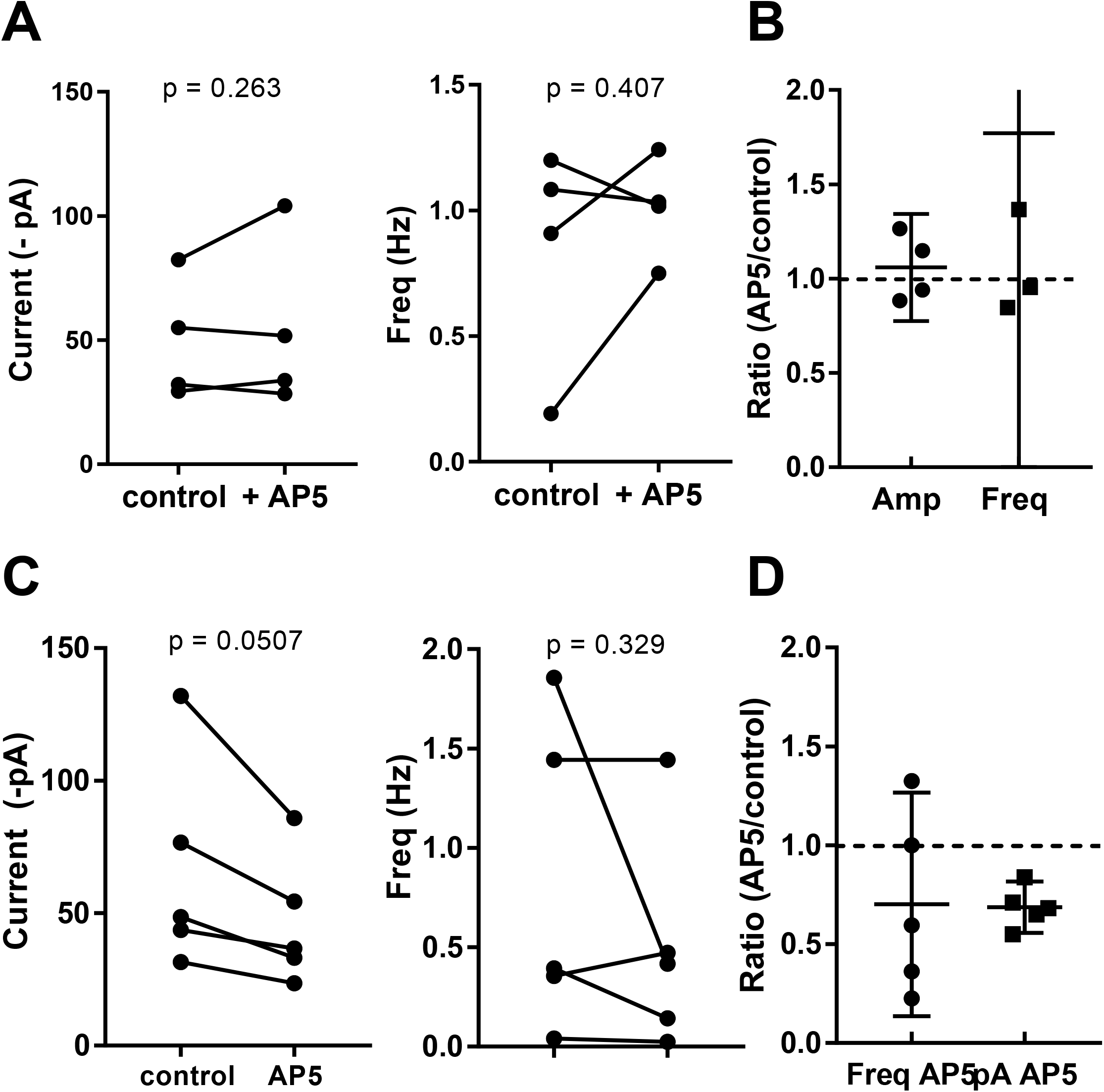
The average peak amplitude (A) and frequency (B) of glycine mIPSCs before (control) and after AP5 (100 μM) exposure when the intracellular solution was altered to include high concentrations of BAPTA. Data points from individual experiments are connected by a line and in B. the proportional change (Ration AP5/control) of mIPSC amplitude from individual experiments is shown. C. The average peak amplitude and frequency of glycine mIPSCs recorded from rat parasaggital spinal cord slices before (control) and after AP5 exposure. Data points from individual experiments are connected by a line and in D, the proportional change (ratio AP5/control) of mIPSC amplitude from individual experiments are shown. The error bars shows the mean and its 95% confidence interval from n = 4-5 cells.

Next, we hypothesised that the lack of effect of AP5 and NMDA on glycine currents was due to species difference, as the original experiments were carried out in cultures derived from rat spinal cord and here we have recorded glycine currents from neurons derived from mouse spinal cord. We prepared parasagittal spinal cord slices from adult Sprague Dawley rats and recorded mIPSCs in the presence of TTX (Figure 4C). The amplitude and frequency of mIPSC recorded were not changed by AP5 (Figure 4C). The mean amplitude in the presence of AP5 was 0.687 times that of control currents (95% confidence interval:0.5571 to 0.8165; P = 0.0507 in two-tailed paired t-test), and the mean frequency was 0.7012 times that of control; confidence interval:0.137 to 1.267; P = 0.329 in two-tailed paired t-test). Note, low mIPSC frequencies measured over the short periods sampled have resulted in the augmented ratio frequency changes.

### Glutamate does not alter recombinant GlyR currents

Finally, we tested the ability of glutamate to enhance currents generated by recombinant glycine receptors expressed in Xenopus laevis oocytes. In the original study, AP5 and glutamate enhanced both α1 and α1β GlyR currents expressed in HEK cells. This finding strongly indicated that the GlyR potentiation observed was due to a direct interaction of glutamate and AP5 with GlyR and not due to an interaction with another receptor or closely associated protein. Whole cell glycine currents were stimulated in oocytes expressing human α1 or α1β glycine receptors with bath applied glycine (10 μM) in the presence and absence of glutamate and blocked by strychnine (Figure 5A-C). Glutamate alone did not stimulate any current (Figure 5A,B) and the mean amplitude of the glycine induced current was unchanged by the presence of glutamate in both glycine receptor subtypes (Figure 5D; GlyRα1: 0.928 of control, confidence interval:0.763 to 1.093; GlyRα1β, 0.927 of control; confidence interval:0.696; adjusted P = 0.755, repeat measure one-way ANOVA).

**Figure 5:**
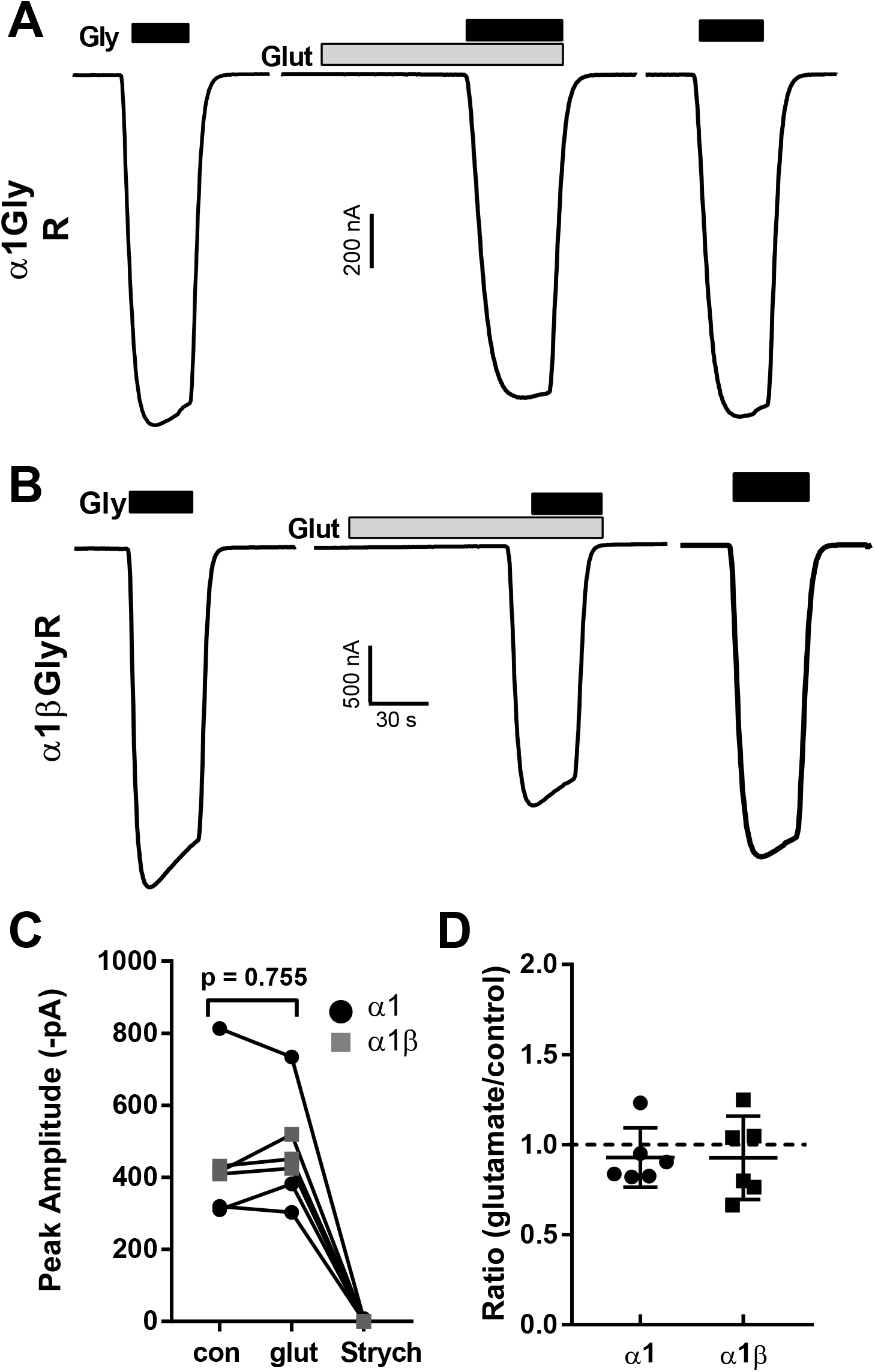
Example of currents induced by extracellular application of glycine (10 μM, black bar) before and after glutamate pre-incubation (100 μM, grey bar) to oocytes expressing α1 (A) or α1β (B) glycine receptors. C. Raw current amplitudes stimulated with 10 μM glycine, after glutamate pre-incubation and then in the presence of strychnine (1 μM). Data points from individual experiments are connected by a line. D. Proportional changes (Ratio glutamate/control) of whole-cell glycine currents from recordings of currents of α1 (circles) or α1β (square) glycine receptors. Data from cells where glutamate was pre-applied for >1 minute before glycine application are combined with cells where glutamate and glycine were directly co-applied. The error bars shows the mean and its 95% confidence interval from a total of n = 6 (3 of each) cells.

## 9. Discussion

We were unable to replicate the finding that AP5, NMDA and glutamate are PAMs of GlyRs. We recorded GlyR mediated currents stimulated by spontaneous and evoked synaptic release of glycine, as well as by exogenous bath-applied glycine. We tested the ability of AP5, NMDA and glutamate to enhance GlyR on mouse, rat and recombinant human GlyRs at low and high glycine concentrations, and confirmed that GlyRs can be potentiated by zinc, suggesting that synaptic release of glycine is not saturating in our cultures. Thus, our failure to reproduce the previously published results cannot be explained by differences in subunit composition, species difference or receptor saturation and suggests that these compounds do not have a direct modulatory effect on GlyRs.

### Methodological differences

We were careful to replicate the key methodical conditions described in Lui et. al. (2010). The slight differences in internal pipette solution were not responsible for the lack of effect observed as confirmed in Figure 4. Similar to the original study, we pre-exposed neurons to AP5/NMDA for >3 minutes before recording mIPSC, eIPSCs or exogenous glycine currents in neurons. In addition, GlyR expressing oocytes exposed to glutamate for >1 minutes before the GlyR current was stimulated were not altered compared to control currents (see Fig. 6).

### Cell selection

We considered the possibility that the GlyR enhancement reported in Lui et. al. (2010) was the result of the enrichment of a receptor subtype that is uncommon in our cell cultures, or the unconscious selection of different neuronal subsets by the two groups. However, given Lui et. al. also observed the effect on GlyRs expressed in HEK cells this possibility is unlikely. In addition, we confirmed that mIPSCs recorded in our spinal cord cultured neurons were enhanced by zinc, a well described PAM of the GlyR (Suwa et al., 2001). Together, these findings indicate that the recorded cell population is not a major source of the difference between this study and the study of Liu et al (2010) and demonstrates that GlyR are not saturated by the synaptic release of glycine in our cultured spinal cord neurons.

### Contaminants

In addition to zinc, there are a host of other compounds that have been reported to strongly directly enhance glycine receptor function, including anaesthetics, cannabinoids and other fatty compounds (Hejazi et al., 2006; Yang et al., 2008), alcohol (Burgos et al., 2015; Lara et al., 2019), intracellular calcium (Fucile et al., 2000), glucose (Breitinger et al., 2015) and others (Huang et al., 2017; Yevenes and Zeilhofer, 2011). This indicates that the glycine receptor likely has multiple PAM binding sites able to unlock the receptor and significantly increase glycine receptor mediated signalling. It is possible that a known or unknown PAM contaminant is responsible for the enhancement observed by Lui et. al. (2010). Given that the enhancement was observed in response to many different compounds, and observed at the level of single channel and in recordings of recombinant GlyRs, the presumptive contaminant could conceivably be part of the vehicle that the drug solutions were made in, as this would have been added to both the drug and control solutions. Alternatively, if glycine and glycine + drug solutions were always delivered from the same drug reservoir, it is also possible that low level contamination of some high affinity positive modulator was present in the drug delivery system and caused these conflicting results. We wash our drug delivery reservoir and tubing regularly with 70% v/v ethanol to prevent this sort of contamination. Finally, glycine itself may be a contaminant of solutions as it is a breakdown product of microbial contaminants. It is conceivable that drug solutions dissolved in water based vehicle that are used over extended periods of time may contain significant concentrations of glycine, especially if stored at room temperature (Hamilton and Myoda, 1974).

Other possibilities considered were that sequentially applied compounds could interact with the glycine receptor to expose and bind to an unusual PAM binding site, or that the compounds used uncovered a strychnine sensitive current that does not originate from the glycine receptor. However given Lui et al. (2010) were able to detect GlyR enhancement in recombinant systems and single channel recordings these possibilities are unlikely.

### Intracellular Calcium Elevations

Intracellular calcium increases are a common feature of neuronal signalling and many reports have shown that various neuronal activators cause intracellular calcium dependent increases in GlyR (Fucile et al., 2000; Kloc et al., 2019; Xu et al., 2000) (Lévi et al., 2008) and GABAR (Stelzer and Wong, 1989) signalling. In essence this means that any process that leads to elevations in intracellular Ca^2+^ will enhance GlyRs. Thus, in addition to contaminants that act directly on GlyR as a PAM, any contaminant that cause an elevation of intracellular Ca2+ could also be the origin of the data reported in Liu et. al. (2010). A recently published paper investigating the NMDA receptor – intracellular Ca^2+^ link to GlyR signalling (Kloc et al., 2019) recorded glycine eIPSCs in the presence and absence of NMDA or D-AP5. Although these experiment were not specifically designed to test the findings reported in Liu et al 2010, this data demonstrates that glycine eIPSCs are not enhanced by NMDA (50 μM) when NMDAR are blocked by the non-competitive antagonist 7-chlorokynurenic acid or when 15 mM EGTA is included in the intracellular pipette solution (Kloc et al., 2019). This data is consistent with our finding that NMDA does not have a direct effect on glycine receptors.

### Main conclusions

This study disputes the previously published finding that AP5, NMDA and glutamate act directly at GlyRs as positive allosteric modulators (Liu et al., 2010) and indicates that the finding that glutamate spill over onto inhibitory synapses interacts directly with GlyR to facilitate neurotransmitter crosstalk and help balance inhibitory and excitatory transmission in the spinal cord and hind brain is false. This changes our understanding of how synapses function and are regulated.

Neurobiological research is particularly susceptible to experimental confounds as the brain is designed to adapt and respond to subtle changes in input and is modulated by compounds that can have exceptionally high affinities (pM - μM) for their biological target. This is significant because very low levels of contaminants can have profound biological effects. As unknown mechanisms of modulation are difficult to control for and brain function is highly dependent on physiological context, the publication of replication studies of key neurobiological findings by independent laboratories is essential to consolidating knowledge and more quickly advance understanding and development in the field of neurobiology.

## Abbreviations

AP5: D-(−)-2-Amino-5-phosphonopentanoic acid
NMDA: N-methyl-D-aspartate
MK 801: (5S,10R)-(+)-5-Methyl-10,11-dihydro-5H-dibenzo[a,d]cyclohepten-5,10-imine
TBOA: threo-β-Benzyloxyaspartic acid
GABA: g-aminobutyric acid
AMPA: α-amino-3-hydroxy-5-methyl-4-isoxazolepropionic acid

## 10. Acknowledgments

Thanks to Bryony Winters, Sherelle Casey and Thomas Harmon for providing critical feedback.

## 11. Authorship contributions

Participated in research design: KA, YO

Conducted experiments: KA, DS

Contributed new reagents or analytic tools: n/a

Performed data analysis: KA, DS

Wrote or contributed to the writing of the manuscript: KA, YO, RV, DS

## 13. Footnotes

KA and YO were supported by the Pain Foundation Ltd. (www.painfoundation.org.au, ABN 87 072 480 123, Registered Charity CFN 21042); RV and DS were supported by the NHMRC (Project Grant APP1144429); and DS holds a Postgraduate Scholarship from the University of Sydney A version of this article has been submitted to the pre-print server bioRxivs (https://www.biorxiv.org/)

